# Accurate MAG reconstruction from complex soil microbiome through combined short- and HiFi long-reads metagenomics

**DOI:** 10.1101/2025.09.12.675765

**Authors:** Carole Belliardo, Nicolas Maurice, Arthur Pere, Samuel Mondy, Alain Franc, Marc Bailly-Bechet, Claire Lemaitre, Riccardo Vicedomini, Jean-Marc Frigerio, Franck Salin, Élodie Belmonte, Marie Gislard, David James Sherman, Pierre Abad, Clémence Frioux, Étienne G.J. Danchin

## Abstract

**Background:** Advances in high-fidelity long-read (HiFi-LR) sequencing technologies have opened new opportunities to explore the microbial genomic diversity of complex environments, such as soils. While short-read (SR) sequencing has enabled broad insights at the gene level, the limited read length constrains the reconstruction of complete genomes. HiFi-LRs, in contrast, improve assembly continuity and completeness, supporting higher-resolution taxonomic and functional annotation. However, the cost and relatively low throughput of HiFi-LR sequencing can limit genome recovery—particularly at the binning stage, where coverage depth is critical. In this study, we assess the benefit of combining HiFi-LR and SR sequencing for genome-resolved characterization of a soil microbiome.

**Results:** We generated metagenomic data for a tunnel-cultivated soil sample using high coverage Illumina SRs as well as a combination of two HiFi-LR sequencing platforms (PacBio Sequel II and PacBio Revio). We found that assemblies generated from pooled HiFi-LR data alone exhibited higher completeness compared to those from ultra-deep SR data. Incorporating SR-derived coverage information for the binning of HiFi-LR contigs further increased both the number and quality of recovered metagenome-assembled genomes (MAGs), with a 24% increase in MAG recovery (313 vs. 252) and lower contamination levels (116 vs. 132 contaminated bins; mean 7.09 vs. 8.07), compared to using HiFi-LR data alone. This approach enabled the recovery of 61 additional MAGs, including 67% of low-abundance and taxonomically diverse lineages such as Archaea, representing 36 novel lineages.

**Conclusion:** Our results demonstrate that integrating HiFi-LR and SR sequencing markedly enhances genome recovery and binning accuracy in a highly diverse environment such as soil. The hybrid approach employed leverages the strengths of both technologies, leading to more contiguous assemblies and enabling the recovery of a broader range of genomes, including low-abundance and taxonomically diverse taxa. While factors such as sequencing depth, cost, and DNA quality remain important considerations, our study provides practical guidance for designing future soil metagenomics projects and underscores the value of adopting long-read technologies for more comprehensive characterization of complex microbial communities.

## Background

Soil is one of the most diverse microbial ecosystems and remains underexplored [1, 2]. It hosts an extraordinary diversity of microorganisms, including bacteria, archaea, fungi, protists, and microscopic animals. The work of Torsvik et al. [3] reports that a single gram of soil may contain up to 10^10^ microbial cells covering diverse taxa, and may harbour up to thousands of different species. Compared to host-associated biomes such as the human gut, soil microbiomes exhibit much greater alpha-diversity and a higher proportion of unknown taxa. Remarkably, just ∼2% of bacterial phylotypes account for nearly 50% of all bacterial sequences in soils worldwide [4]. Comprehensive characterization of soil microbiomes, especially rare species, is essential for advancing our understanding of biogeochemical cycles [5] and developing sustainable agricultural practices [6]. Microbial genomes encode the genes responsible for the metabolic capacities and ecological functions that shape soil processes, including interactions, catalysis of biochemical reactions, and contributions to ecosystem functioning [7]. Thus, uncovering the diversity of soil microbial genomes is crucial for harnessing microbial resources and promoting soil health and sustainability [6].

Various approaches have been developed to characterise soil microbial communities. Although culturomics has provided valuable insights, many soil species remain uncultivable under laboratory conditions [8]. Consequently, metagenomics and other culture-independent omics approaches have become the primary strategies for exploring the taxonomic diversity and functional potential of soil microbiomes. Shotgun metagenomics allows direct DNA sequencing from complex microbial communities without prior cultivation. This approach can provide a genomic snapshot of the microbial community in a given soil sample, even if dark matter, constituted by rare species, remains difficult to capture. Metagenomics has revolutionised our ability to investigate the diversity and functions of uncultured organisms, providing unprecedented insights into the genetic diversity, functional potential, and ecological roles of microbiota [9]. *De novo* metagenomic assembly allows the reconstruction of genomes species, including uncultured taxa, and the prediction of the encoded proteins [10, 11]. However, the reconstruction of high-quality genomes from the complex mixture of sequenced reads remains very limited in soil, due to the extremely high microbial diversity, uneven species abundance, and the presence of closely related strains, which together complicate assembly and binning processes. Consequently, only a few whole genomes are usually obtained, with many unassembled reads [12].

Most metagenomic projects conducted over the past decade have relied on short-read (SR) sequencing technologies, generating high-throughput datasets typically composed of 2 × 50–150-base-pair fragments. Consequently, SR-based metagenomic studies still dominate public repositories such as the JGI IMG/M [13] and EBI MGnify [14] databases in terms of available sequence data. Despite their high accuracy and throughput, SR remains limited to face the challenge of reconstructing genomes containing multicopy genes or repeat-rich regions, which are inherently difficult to resolve with short sequences, creating ambiguous paths in the assembly graph and leading to fragmented assemblies [15]. In the 2022 release of the JGI IMG/M platform, only 10% of SR-derived contigs exceeded one kilobase in length, illustrating the extent of assembly fragmentation. Moreover, inconsistencies in protein taxonomic assignments within contigs suggest the presence of chimeric assemblies [11].

Increased read length in metagenomic studies enhances contig lengths by providing long-range information, which allows better resolution of repetitive sequences. In recent years, Oxford Nanopore Technologies (ONT) [16, 17] and Pacific Biosciences (PacBio) [18] platforms have significantly improved read accuracy, length, and throughput. While the first generations of LR sequencing technologies were characterised by high error rates, which led to the recommendation of combining them with SR for error correction during the assembly step [19], recent technologies provide high-fidelity (HiFi) sequences, with error rates that are similarly low to those of SR [20, 21]. To leverage the complementary strengths of SR and LR technologies, several hybrid metagenomic strategies have been developed. For assembly, tools such as OPERA-MS [16] use LR data primarily to scaffold contigs initially assembled from SR reads. More recently, HyLight integrated HiFi-LR and SR from low-depth metagenomes [22] and MetaPlatanus [23] implemented mutual-support strategies or overlap-graph approaches to achieve strain-level resolution. Integrated workflows such as MUFFIN [24] further combine hybrid assembly with differential coverage binning to recover high-quality metagenome-assembled genomes (MAGs). Recent benchmarking studies additionally demonstrated that multi-sample hybrid binning strategies outperform other approaches for recovering near-complete strains and detecting antimicrobial resistance genes [25].

Although several studies have explored hybrid approaches for metagenomic assembly and binning, a strategy relying exclusively on LR data for assembly while using SR data exclusively for binning remains unexplored. In this study, we assessed the added value of HiFi-LR sequencing compared to SR sequencing for metagenomic assembly in complex environments such as soil. We generated and analysed PacBio HiFi-LRs, Illumina SRs, and metabarcoding data from a cultivated soil sample, comparing contigs and MAG recovery for each individual technology and for both combined. Our results demonstrate that a hybrid strategy, combining the higher contiguity of HiFi-LR assemblies with the greater sequencing depth of SR data, improves genome reconstruction. Incorporating SR-derived coverage into HiFi-LR binning led to an increase in MAG recovery (+ 24%), and reduced the global contamination levels compared to HiFi-LR data alone. The gain is particularly marked for low-abundance taxa. Comparison with microbial diversity estimates from metabarcoding data confirmed that this hybrid approach captures a broader spectrum of the soil microbiome.

## Methods

### Sampling

We collected a soil sample between two rows of salad crops in a tunnel farming setup in Lambesc, southern France. Rapid freezing, followed by subsequent freeze-drying procedures at -40°C, immediately preserved the soil sample. These preservation and preparation steps were meticulously executed at the Conservatoire de Ressources Génétiques (CRG), hosted within the Genosol platform in Dijon, France, to maintain the molecular integrity of the samples in readiness for subsequent analytical procedures.

### DNA extraction

Total genomic DNA was extracted using modified IS0-11063 protocols for DNA extraction and Nucleospin soil for DNA purification [26]. From 1g of dry equivalent soil (freeze-dried), DNA was extracted by mechanical lysis using FastPrep 5G (MP Bio) coupled with cell lysis with denaturing detergent (lysis buffer: EDTA, Tris pH 8.0, NaCl and SDS 2%). Deproteinisation consisted of precipitating the proteins with a high salt concentration (KAc 5M) followed by high-speed centrifugation to separate the DNA (supernatant) and proteins (pellet). DNA has been precipitated with isopropanol (v/v) and washed with ethanol 70%. DNA was quantified with NanoDrop technology with a concentration of 114.7 ng/*µ*L (260/280 ratio of 1.84, 260/230 ratio of 1.5). As Nanodrop could overestimate the DNA content, the QuBit method, a more reliable method for DNA quantification, was used and yielded a dosage of 50.5 ng/*µ*L (total DNA quantity 2.62 *µ*g in a volume of ca. 50*µ*L). Notably, the sample exhibited a straightforward appearance, denoting its suitability for further investigation.

### Metagenomic and metabarcoding sequencing and raw data processing

#### Long-read metagenomic sequencing

Extracted soil DNA was sequenced on the PacBio platforms using the HiFi sequencing mode. One complete SMRT cell was run on both the Sequel II and Revio platforms, generating two datasets. For the Sequel II sequencing, high-molecular-weight DNA with an average fragment size of ∼8 Kb was used to prepare an unamplified, non-multiplexed HiFi library with the Template Prep Kit v2.0, following the PacBio metagenomic shotgun protocol. Sequencing was carried out on a SMRT Cell 8M over 30 h. For the Revio sequencing, the HiFi library was prepared with ∼8 Kb not-sheared native DNA, using the Template Prep Kit 3.0. The SMRTBell library was sequenced on a SMRTCell 25M over 24 h with the Revio sequencing plate reagents. For both Sequel II and Revio datasets, adapter removal, read quality filtering (minimum Q20), and generation of circular consensus sequencing (CCS) reads were performed using dedicated PacBio pipelines. HiFi-LR reads of both platforms were pooled to increase sequencing depth.

#### Short-read metagenomic sequencing

DNA from the same soil sample underwent a second extraction and was sequenced at ultra-deep coverage on a full NextSeq 2000 Illumina P3 flow cell.

#### 16S rRNA gene amplicon sequencing

16S rRNA gene amplicon sequencing data were generated from the same soil sample through targeted sequencing of the V3-V4 region of the 16S rRNA gene using primers 341F_UDI and 785R_UDI on the Illumina NextSeq 2000 platform, producing 2 × 300bp reads. Library preparation and sequencing were conducted with the protocol described by [27]. Based on the selected primers, we expected to sequence approximately 444 bp of the 16S rRNA gene, with an expected overlap of approximately 156 bp between R1 and R2. Paired-end reads were merged using FLASH v0.23.4 [28] with flexible overlap settings (--min-overlap 10, --max-overlap 301) to reconstruct full-length amplicons, even with region size variability. Primer sequences were trimmed using Cutadapt v4.9 [29] with parameters optimised for mismatch handling (--error-rate 0.1, --match-read-wildcards, --overlap). Quality filtering was performed using Fastp v0.23.4 [30], with the following parameters: a Phred quality score threshold of 30 (--qualified_quality_phred 30), a minimum sequence length of 230 bp (--length_required 230), and an average quality of 30 (--average_qual). Additional trimming at the 5’ and 3’ ends was done using a sliding window approach (--cut_window_size 4, --cut_mean_quality 30) with low-quality bases removed (--cut_front, --cut_right). Adaptor trimming was disabled (--disable_adapter_trimming) to avoid interference with prior primer removal steps. Chimeric sequences were detected and removed using Vsearch v2.9.1 [31] in *de novo* mode (--uchime3_denovo).

### Bioinformatic analyses

A summary of the bioinformatic pipeline and metagenomic processing is available in Supp. Fig. 1. After data acquisition, we performed quality control on each dataset using FastQC software version 0.12.0 [32] with default parameters. The results were merged using MultiQC software version 1.25.1 [33] in default mode.

#### Assembly

Paired-end metagenomic SR were assembled using MEGAHIT v1.2.9 [34] with the ‘--presets meta-large’ parameter. HiFi-LR were assembled using MetaMDBG v.0.3 with default parameters [35].

#### Binning

We used three binning strategies according to the sequencing data used to perform alignments against contigs (Supp. Fig. 1). Sequence features, such as GC content, tetranucleotide frequency, and coverage as a proxy for abundance, were used for clustering contigs that are supposed to belong to the same organism in the bin, toward the reconstruction of MAGs.

- *Binning strategy* 1: short-read assembly with short read mapping for contig binning (SR-SR).
- *Binning strategy* 2: long-read assembly with long read mapping for contig binning (LR-LR).
- *Binning strategy* 3: long-read assembly with short read mapping for contig binning (Hybrid SR-LR strategy).

In binning strategies 1 and 3, short reads were mapped on contigs using BWA-MEM2 v.2.2 [36] with default parameters. In Binning strategy 2, long reads were mapped with Minimap2 v.2.26 [37] with the parameters -x map-hifi -a --sam-hit-only --secondary=no.

In all the binning strategies, resulting alignments were used to compute contig coverage, which served as a proxy for estimating species abundance. Mapping outputs were processed using Samtools v.1.10 [38]. Then, in all three strategies, contigs were binned using MetaBAT2 v.2.15 [39] in default mode, and SemiBin v.1.5.0 [40] with parameters ‘--environment soil’ and ‘--sequencing-type=long_read’ for long-read processing (Binning strategies 2 and 3). The obtained results were merged and dereplicated using DAStool v.1.1.6 [41].

#### Bin and Metagenome-aAssembled Genome (MAG) evaluation

Gene prediction was performed using Prodigal v2.6.3 [42], and the completeness and contamination of bins were assessed using CheckM2 v1.0.1 [43]. Those analyses were performed using a previously published workflow [44]. We required at least 50% completeness and less than 10% contamination, without any filter on the number of contigs, for MAGs and taxonomically assigned bins. Dereplicated bins were classified based on their completeness and contamination levels, as described in the MIMAGs guidelines [45]. Bins with 100% completeness and ≤5% contamination were considered near-complete MAGs. Bins reaching the ≥90% completeness criterion were labelled high-quality (HQ) MAGs. Those comprising between ≤90% and ≥50% completeness were considered medium-quality (MQ) MAGs. Bins below these thresholds were considered low-quality (LQ) MAGs, while those over 10% of contamination were classified as ‘*contaminated bins*’. Metrics regarding assembled and non-assembled reads (e.g., k-mers, GC content, read size) and assembly and binning quality were obtained using Mapler v2.0.0 [46].

##### Subsampling of short-reads for coverage calculation at the binning step

To evaluate the impact of short-read sequencing depth on contig binning efficiency, additional subsampling analyses were performed on the Hybrid SR-LR strategy (binning strategy 3). SR datasets were randomly subsampled to 10% and 50% of the initial sequencing depth using SeqKit v.2.13.0 [47] with the command seqkit sample2 -p and default parameters. The resulting subsets of short reads were independently mapped against LR-sq+rv contigs using BWA-MEM2 v.2.2, and contigs were subsequently binned following the same workflow described above using MetaBAT2, SemiBin, and DAS Tool.

#### Taxonomic classification

To characterise the taxonomic composition of MAGs, contigs, and reads from each dataset, two complementary taxonomic assignment strategies were applied. For MAGs with more than 50% completeness, taxonomic classification was confirmed using GTDB-tk v.2.1.1 [48] with database version 214, providing robust information based on multigene phylogenetic placement [49].

Additionally, both Illumina 16S rRNA amplicon sequences and PacBio metagenomic contigs as well as reads underwent a common taxonomic assignment strategy to facilitate their comparison. Reads (metabarcoding and LR metagenomics) and contigs (LR and SR metagenomics) were assigned a taxon using MMseqs2 [50] “easy-search” module against the SILVA 138.2 SSURef NR99 database of curated 16S rRNA sequences as the reference. To ensure high-confidence assignments, homology searches were performed using a stringent threshold of 97% sequence identity and a minimum alignment length of 100 bp (parameters --min-seq-id 0.97 and --min-aln-len 100, respectively). Coverage was set to improve sensitivity in detecting partial sequence matches of fragmented SR and LR reads and contigs by including overlapping hits (--cov-mode 2). Alignment coverage was analysed using BEDtools [51], calculating the proportion of each reference sequence covered by aligned reads. Species with 16S sequences from SILVA that were covered on more than 97% of their length were retained for further study. Eukaryotic hits against the SILVA database [52] were excluded to ensure consistency in taxonomic inference. The taxonomic lineage was used to increment counts in an abundance matrix for each taxonomic rank from species to domain. These matrices underwent downstream diversity analyses, which were performed using custom Jupyter Notebooks. Phylogenetic visualisations were generated using the Interactive Tree Of Life (iTOL) v7 [53].

## Results

### HiFi-LR sequencing delivers more comprehensive and continuous metagenome reconstruction

To compare shotgun metagenomic assembly performance of short-read and long-read sequencing technologies, we sequenced a soil sample with Illumina SR and PacBio HiFi-LR (Sequel II and Revio) platforms and assembled the resulting datasets. Our data show that HiFi-LR sequencing, particularly with the Revio platform, yields more comprehensive reconstructions of microbial genomes with enhanced continuity and more complete contigs.

Ultra-deep sequencing with paired-end Illumina technology generated 903.3 million reads of 2 × 150 base pairs in length. Combining PacBio HiFi-LRsequencing using a full SMRTcell on Sequel II and another on Revio platforms yielded a lower total of 11.2 million reads, but with an average length of 7.25 kb, and a maximum length of 40 kb (Table 1). The read length distribution of our metagenome sequencing aligns with DNA fragment size estimates for soil samples, mirroring findings from human gut microbiome studies [18].

**Table 1:**
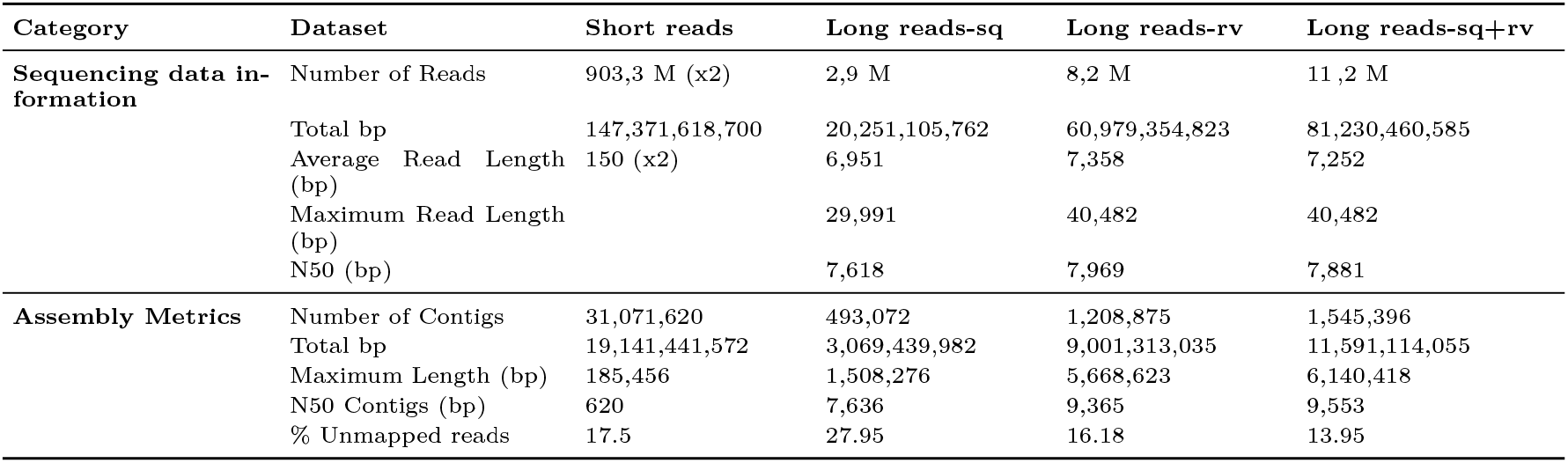
Summary of Illumina Next-seq 2000 short reads and PacBio long reads from Sequel II (sq), Revio (rv) and pooled Sequel and Revio (sq+rv). *”M” = Millions, “G”=Giga, “bp” = base pairs, “sq” = sequel, “rv” = revio.

| Category | Dataset | Short reads | Long reads-sq | Long reads-rv | Long reads-sq+rv |
| --- | --- | --- | --- | --- | --- |
| **Sequencing data information** | Number of Reads | 903,3 M (x2) | 2,9 M | 8,2 M | 11 ,2 M |
|  | Total bp | 147,371,618,700 | 20,251,105,762 | 60,979,354,823 | 81,230,460,585 |
|  | Average Read Length (bp) | 150 (x2) | 6,951 | 7,358 | 7,252 |
|  | Maximum Read Length (bp) |  | 29,991 | 40,482 | 40,482 |
|  | N50 (bp) |  | 7,618 | 7,969 | 7,881 |
| **Assembly Metrics** | Number of Contigs | 31,071,620 | 493,072 | 1,208,875 | 1,545,396 |
|  | Total bp | 19,141,441,572 | 3,069,439,982 | 9,001,313,035 | 11,591,114,055 |
|  | Maximum Length (bp) | 185,456 | 1,508,276 | 5,668,623 | 6,140,418 |
|  | N50 Contigs (bp) | 620 | 7,636 | 9,365 | 9,553 |
|  | % Unmapped reads | 17.5 | 27.95 | 16.18 | 13.95 |

#### HiFi-LR sequences result in more comprehensive metagenome assembly

Reads were assembled using technology-specific pipelines to ultimately reconstruct the genomes of soil microorganisms. A first observation is that, before assembly, raw HiFi read lengths substantially surpass the lengths of contigs assembled from SR, as previously observed in publicly available soil metagenomes, where the average size of SR contigs was less than 1kb [11]. Our dataset confirms these observations, with SR contigs having an N50 of 620 bp and a largest contig size of 192 kb, whereas unassembled HiFi-LRs have an N50 of 7,618 bp. The assembly of PacBio HiFi-LRs further increases this length difference. Sequel II and Revio long reads were both assembled separately, and also pooled (sq+rv) as a third HiFi-LR dataset, which was assembled to compare the effect of technologies and depth (Sequel II < Revio < merged Sequel + Revio) on assembly quality. Revio generated three times more base pairs than Sequel II (Table 1), a trend that persisted at the assembly level, with Revio assemblies containing three times more assembled base pairs in total (Table 1).

Four contigs obtained from Sequel II reads were longer than 1 Mb, with the longest being 1.5Mb in length. In contrast, assembly of Revio reads yielded 139 contigs of at least 1 Mb, and 87 longer than the longest Sequel II contig. The longest contig obtained with Revio was 5.7 Mb long, demonstrating the capacity of this technology to deliver HiFi-LRs with higher coverage useful for resolving microbial genomes from complex environments. Although Illumina yielded nearly twice the total number of base pairs compared to PacBio at both the read and contig levels, the short-read assembly remained highly fragmented, with an N50 of only 620 bp and a maximum contig length of 185 kb (Table 1, Fig. 1). The Revio platform, by generating more HiFi-LRs, enables the assembly of contigs longer than those generated not only from Illumina data but also from Sequel II data.

**Figure 1:**
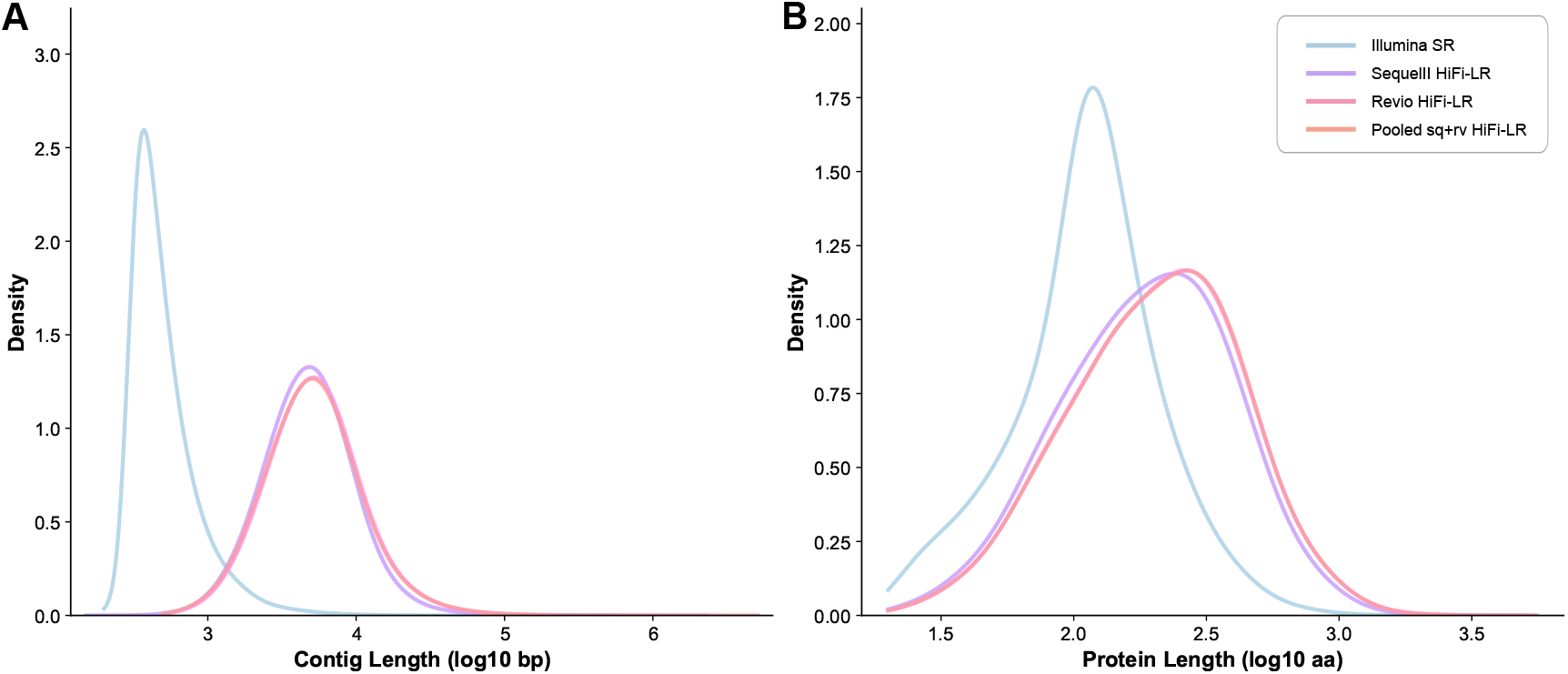
**A)** Contig size and **B)** protein length distribution (log scale) for short reads, SequelII and Revio as well as pooled sq+rv HiFi-LRs.

The improvement in contig lengths generated from long reads is expected to positively impact the downstream completeness of gene predictions. *De novo* gene prediction from long-read assemblies yielded approximately 18 million proteins classified as complete, representing 80% of all predictions, compared to fewer than 5 million (12% of all predicted proteins) from SR contigs. Furthermore, the length distributions of predicted proteins are consistent with those reported by Never and colleagues [54] for high-quality predicted proteomes (Supp Fig. 1B). Indeed, in this work, the authors showed that protein length distribution is a reliable indicator of genome assembly quality, with conserved protein lengths typically mostly ranging from 236 to 353 amino acids (726 to 1059 base pairs) across different domains of the tree of life (archaea: 242 amino acids, bacteria: 270 amino acids, eukaryotes: 353 amino acids). This increased protein prediction accuracy is observed in both the LR contigs and the predictions from raw LR reads.

Although HiFi-LRs generated by the Revio platform yielded better assembly metrics and gene predictions than short reads, combining the two long-read datasets (Sequel II + Revio, sq+rv) further enhanced assembly metrics. This suggests a direct positive effect of HiFi-LR sequencing depth on the assembly statistics. While the contig length is only slightly improved (Fig. 1, Supp. Fig. 2), the number of contigs and total base pairs incorporated in contigs for the pooled sq+rv dataset have significantly increased (Table 1). These results confirm the importance of sequencing depth in reconstructing DNA sequences from complex environments [55].

#### Unmapped reads indicate limitations in coverage and assembly, mainly in low-abundance species

Unmapped reads are those that fail to align with the assembly either because they contain sequencing errors or are not represented in any contig. Characterising unmapped reads is crucial for assessing sequencing completeness and identifying genomic regions (or even species) that are challenging to assemble. In metagenomics, those unmapped reads often originate from low-abundant populations, leading to undetected species or highly fragmented bins.

Here, mapping statistics revealed significant differences between sequencing platforms. When Illumina SR exhibited an unmapped-read rate of 17.5% on their associated contigs, PacBio HiFi-LR demonstrated unmapped-read rates of 28%, 16% and 14% for contigs generated from Sequel II, Revio or both technologies pooled, respectively (Table 1, Fig. 1A). On the PacBio datasets, the higher sequencing depth yielded a significantly lower proportion of unmapped reads (p-value <10^−5^, Z >100 for all comparisons with a two-sample proportion Z-test), with a small to medium effect size, increasing with the difference in sequencing depth (cohen’s h: 0.06 between pooled and Revio, 0.29 between Revio and Sequel II, and 0.35 between pooled and Sequel II). Thus, the higher number of generated contigs and incorporated base pairs increased the diversity of genomic regions captured by the assembly.

Further comparisons of reads (Supp. Fig. 3) highlighted that unassembled long reads (sq+rv) exhibited slightly shorter lengths than those participating in the assembly, with a median length of 6.3 and 7.0 kb, respectively; the distributions in the two groups differing significantly (Mann-Whitney, P < 2.2e^-16^, two-tailed), but with a small effect (rank-biserial correlation, r=0.15). Unassembled reads also displayed a lower GC content than the other reads, with a median of 63.6% compared to a median of 65.8% (Mann-Whitney, P < 2.2e^-16^, two-tailed, rank-biserial correlation: r=0.17). Additionally, their abundance (as calculated by the median of their k-mer occurrences within the full read dataset) was significantly lower, with a median of 1, compared to 3 for the assembled reads (Mann-Whitney, P < 2.2e^-16^ two-tailed), and with a strong effect (rank-biserial correlation, r=0.73).

It is worth noting that, with an average length of 6.7 kb, the unassembled reads could still contain full-length gene sequences. These sequences could uncover genomic fragments and functions, possibly from low-abundance species still uncaptured in assembly-based analyses. These findings suggest that, due to sequencing depth limitations, long-read assemblies still miss a significant portion of the metagenomic diversity, resulting in a substantial number of species left unassembled. This underscores the importance of integrating complementary strategies to enhance the recovery of genomes from complex soil communities.

Finally, to assess overlap between captured biodiversity across technologies, short reads were mapped on long-read assemblies. We observed that 21% of SR remained unmapped; this proportion increased to 62% under stringent mapping parameters (identity >90% and alignment length ≥130 bp), highlighting partial but incomplete overlap in captured biodiversity between both sequencing technologies.

### Using SR coverage improves binning of HiFi-LR contigs

We then compared the number and completeness of genomes reconstructed after binning for both SR and HiFi-LR datasets, considering different mapping strategies prior to binning: (i) mapping Illumina SR to the short-read contigs (SR-SR), (ii) HiFi-LR to the HiFi-LR contigs (LR-LR), and (iii) Illumina SR to the HiFi-LR contigs (SR-LR). This resulted in three distinct binning workflows (Supp. Fig. 1). Since the pooled sq+rv HiFi-LR from both Revio and Sequel II technologies yielded the best assembly results in terms of contig length distribution and lower % of unmapped reads, we consider only this dataset as representative of the HiFi-LR technology from now on.

#### Comparative analysis of bins and MAGs from Illumina SR and PacBio HiFi-LR assemblies obtained through self-mapping coverage

We first considered binning the HiFi-LR assembly using the mapping of the corresponding long reads to the contigs (Fig. 2A). Such binning led to reduced unbinned contig length (Fig. 2B) and generated more than twice the number of bins (2,587 vs 1,187) and nearly twice as many MAGs (252 vs 145) compared to binning the SR contigs with short-read mapping (Table 2, Fig. 2C). This improvement was also reflected by longer contigs (up to 6.7 Mbp vs. 192 Kbp), greater bin lengths after dereplication, and a substantially higher N50 (487 Kbp vs. 9 Kbp), indicating improved contiguity (Table 2). Regarding MAGs, the HiFi-LR assembly produced substantially more high- and medium-quality MAGs (75 HQ and 169 MQ vs. 16 HQ and 128 MQ for SR), as well as a greater number of low-quality MAGs (176 vs. 51). Notably, HiFi-LR assemblies also generated significantly more near-complete MAGs (8 vs none), highlighting their potential for near-complete genome reconstruction, even in very complex microbiomes (Fig. 2C).

**Table 2:** Summary of bins from Illumina and PacBio sequencing obtained using self-coverage or SR coverage for binning of contigs. “Raw bins” correspond to the total number of bins generated independently by MetaBAT2 and SemiBin. “Non-redundant bins” refer to the set of bins retained after refinement and scoring with DAS Tool, which integrates results from multiple binning tools and selects non-redundant bins. MAGs correspond to metagenome-assembled genomes as defined by CheckM criteria (≥50% completeness and ≤10% contamination). **’ M’ = Millions, “G”=Giga, “bp” = base pairs, “Pcb_sq_rv” = contigs from coassembly of sequel and revio data. Note: DAS Tool was used to integrate bins generated by MetaBAT2 and SemiBin. Dereplication refers to the selection of non-redundant bins based on quality scores

| **Binning Strategy**<br>**Metric** | **Processing Step** | **Self-coverage**<br>**Illumina** | **Pcb_sq_rv** | **SR-coverage**<br>**Pcb_sq_rv** |
| --- | --- | --- | --- | --- |
| Number of bins | Raw bins | 1 187 | 2 587 | 2 518 |
|  | Non-redundant bins | 258 | 560 | 627 |
|  | MAGs | 145 | 252 | 313 |
| Binned contigs (all bins) | Non-redundant bins | 146,518 | 83,327 | 113,232 |
|  | MAGs | 68,669 | 20,889 | 31,384 |
| Binned base pairs (Mbp) | Non-redundant bins | 987 | 1 990 | 2 274 |
|  | MAGs | 548 | 995 | 1 196 |
| Max. contig length (Kp) | Non-redundant bins | 192 | 6 697 | 6 697 |
|  | MAGs | 186 | 6 140 | 6 697 |
| N50 (Kb) | Non-redundant bins | 9 | 487 | 429 |
|  | MAGs | 12 | 907 | 750 |

**Figure 2:**
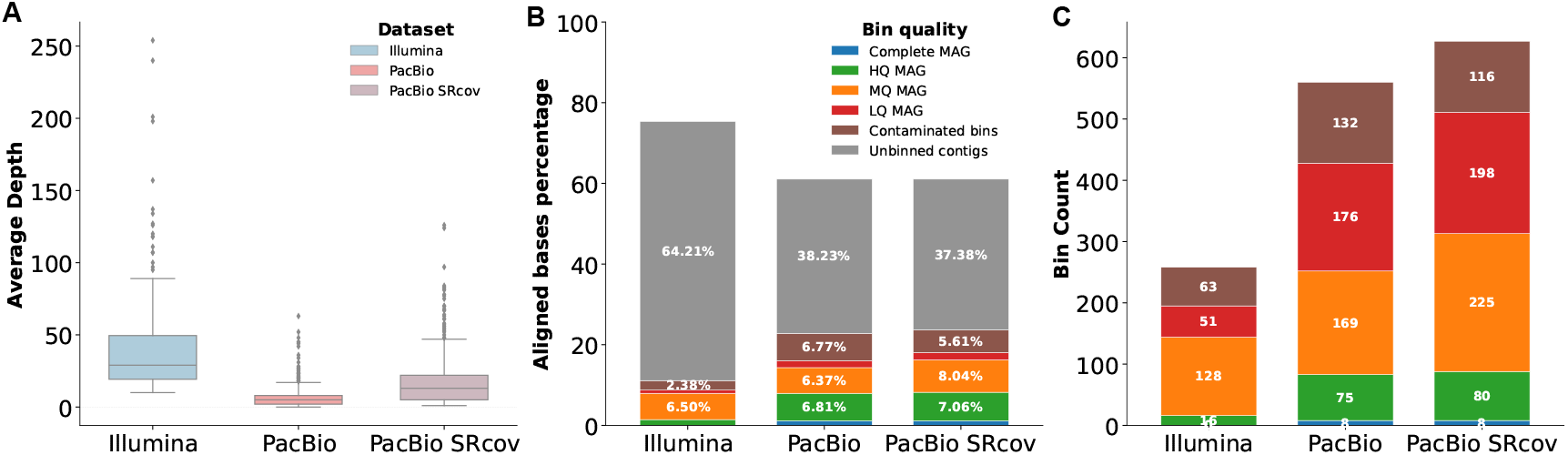
**A)** Average coverage of SR and HiFi-LR contigs after mapping of their respective (SR and HiFi-LR) reads, along with the average coverage of Hifi-LR contigs after SR alignment (PacBio SRcov). **B)** Aligned bases percentage (length of read segments aligned to the assembly divided by total reads length, grouped by bin quality). **C)** Number of bins and their quality classification. SR and HiFI-LR datasets were binned using self-mapping of their respective reads, while “PacBioSRcov” refers to the PacBio assembly binned using SR data. Bin quality categories include Complete, High Quality (HQ), Medium Quality (MQ), Low Quality (LQ), and Contaminated (>10% contamination).

However, these better assembly metrics came with an increase in contamination, as evidenced by a larger number of contaminated bins (>10% contamination estimates) in the HiFi-LR dataset (132 vs. 63 bins classified as contaminated). This pitfall is likely related to the insufficient coverage information in HiFi-LR bins, due to relatively limited sequencing depth (Table 1) and resulting in relatively homogeneous read abundance when mapped to contigs (Fig. 2A,C). Specifically, HiFi-LR self-mapping resulted in low coverage depths on contigs (mean: 3X; range: 0–310X, Fig 2A) and on MAGs (mean: 6X; range: 0–63X), limiting the resolution of the binning process.

In contrast, the depth of SR sequencing led to broader coverage estimation. Self-mapping SRs on their corresponding contigs yielded substantially higher coverage, with mean depths of 33X on SR-assembled contigs (range: 5–17,156X, Fig. 2A) and 40X on MAGs (range: 10–254X). Notably, more than half of Illumina reads aligned stringently to HiFi-LR-assembled contigs, with coverage depths reaching up to 27,323X (for bins mean: 9X; for MAGs mean: 16X range: 1–126X). These results suggest that the limited depth of HiFi-LR self-mapping undermines binning accuracy and results in an increase in contaminated bins (Fig. 2C). Consequently, we hypothesised that incorporating SR coverage into the HiFi-LR binning process could improve bin quality by enhancing binning resolution and reducing contamination.

#### Comparative analysis of genome bins from PacBio HiFi-LR assemblies using self-mapping versus SR coverage mapping

We therefore designed a hybrid binning strategy, using coverage information from Illumina SR mapping on contigs assembled from pooled sq+rv PacBio HiFi-LR and assessed its benefit. Although the raw total number of produced bins was slightly lower (2,587 bins for self LR-self coverage vs. 2,518 for SR coverage), SR coverage of HiFi-LR contigs generated more unique bins. Indeed, dereplication yielded 627 non-redundant bins using SR coverage, representing a more than 10% increase over the 560 bins obtained from self-coverage, which reflects improved resolution of overlapping or ambiguous contigs in the complex samples. This strategy yielded 313 reconstructed MAGs (+24%) compared to. 252 obtained using LR self-coverage (Table 2, Fig. 2C, Supp. Fig. 4, Supp. Fig. 5).

Improvement is observed for MAGs, where an additional 30% of contigs participate in high-quality MAGs (LR mapping: 20,889 vs. SR: 31,384, Fig 2B). As expected, those contigs have a reduced read coverage compared to the ones shared in the two binning strategies (Supp. Fig. 6, Mann–Whitney U test, p = 6.56 × 10^−135^). Therefore, SR coverage allowed for the incorporation of more base pairs (2,274 Mbp vs. 1,990 Mbp with HiFi-LR self-coverage) in bins, and within MAG sequence recovery (1,196 Mbp vs. 995 Mbp). However, bins obtained from SR coverage have a slightly reduced median contiguity (MAG N50 decreased from 907.6 Kbp to 750.9 Kbp). The number of near-complete MAGs remained stable (8), and the number of high-quality MAGs increased slightly from 75 to 80. In parallel, the number of medium-quality MAGs increased from 169 to 225, and low-quality MAGs increased (176 vs 198). These results suggest that the higher coverage provided by SRs allowed more species to be represented as MAGs, although a substantial proportion remained fragmented. The number of contaminated bins decreased substantially with SR coverage (132 vs 116), suggesting better bin purity with the increase of coverage (Fig. 2A).

If direct comparison of genetic diversity between bins is difficult for assessing genome reconstruction quality in uncultivated species, metrics such as completeness, contamination, and MAG taxonomic assignment can aid in evaluation. The increase in contig recruitment driven by SR coverage enhances the recovery of additional MAGS, while exerting only a marginal effect on the completeness of MAG already reconstructed LR coverage (mean completeness estimates of 62.71 HiFi-LR and 63.05 for SR coverage, respectively, Supp. Fig. 4 and 5). In contrast, this coverage improves genome discrimination by reducing contamination levels (mean contamination estimates of 8.07 HiFi-LR and 7.09 SR coverage, respectively).

This improvement was contingent on sufficiently deep short-read coverage. When subsampling the SR dataset to 50%, we obtained 45 additional MQ MAGs compared to self-coverage, while only one HQ MAG present with self-coverage was lost. However, at 10% SR subsampling, binning quality markedly declined, with fewer MAGs recovered at all quality thresholds than with self-coverage (Supp. Fig. 7).

#### Taxonomic analysis of MAGs

We further compared the MAGs obtained with each of the three binning strategies by exploring their taxonomic assignments. A total of 34 GTDB lineages were identified as being specific to the SR-covered HiFi-LR MAGs dataset. They were not recovered in SR contigs nor HiFi-LR self-coverage binning (Fig. 3). Recovered HiFi-LR-SR MAGs spanned over 30 genera or families, of which 20 were also retrieved in Illumina MAGs, supporting the presence of these lineages in the original samples. Although potentially representing binning artefacts, the remaining 10 taxa specific to SR coverage may include additional lineages uncovered by improved SR-driven binning. Only seven HiFi-LR-self coverage genera were absent in MAGs produced with the SR coverage, including five nonetheless recovered in Illumina-based MAGs (Fig. 3A). Moreover, with SR coverage, some contigs previously clustered in the same bin were redistributed across different MAGs. The low coverage from HiFi-LR mapping originally resulted in the collapse of strain or species-level genomes into a unique MAG.

**Figure 3:**
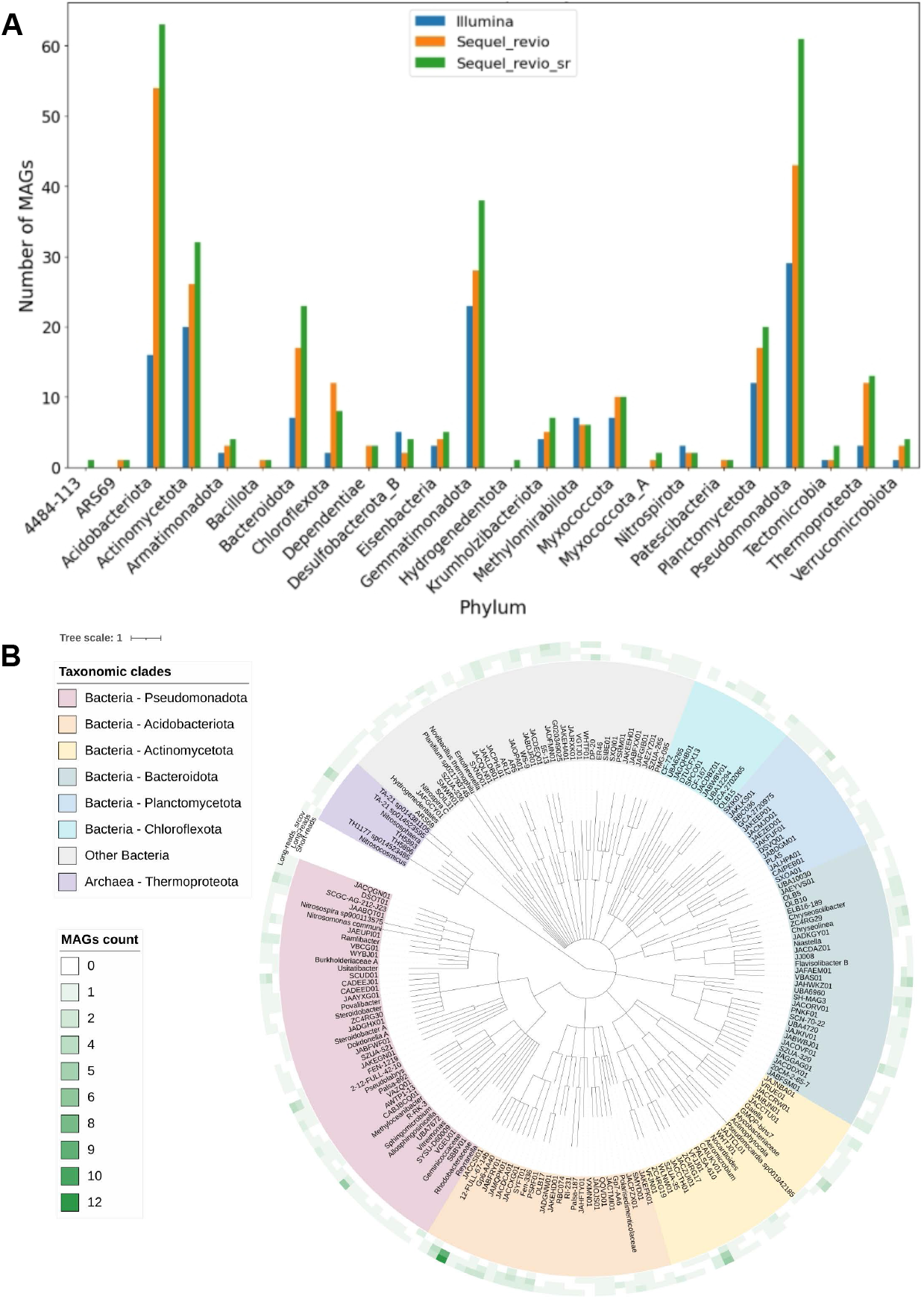
**A)** Taxonomic distribution of metagenome-assembled genomes (MAGs) by phylum across three sequencing and binning strategies, where SR Illumina data are represented in blue, Self-mapped HiFi-LR in orange and HiFi-LR contigs binning with SR coverage in green. **B)** Phylogenetic tree of taxonomic annotations for MAGs, generated using GTDB-tk [48]. The occurrence of each taxon across datasets is visualised as a heatmap. The interactive tree is available at: https://itol.embl.de/export/17616122570468971737508212.

Phylogenetic analysis confirmed that HiFi-LR contigs binned with SR yield the widest taxonomic spectrum (Fig. 3B), recovering previously unbinned clades, such as the bacterial genus SXQI01 from the Verrucomicrobiota phylum. The 48 SR-specific MAGs displayed broad taxonomic diversity, encompassing 17 phyla, and 32 orders. Taxonomic resolution remained high up to the family level (95.8%), while genus- and species-level assignments were obtained for 68.8% and 6.3% of MAGs, respectively. Moreover, the increased number of MAGs across bacterial phyla, such as Planctomycetota and Acidobacteriota, underscores the effectiveness of the hybrid binning strategy in recovering underrepresented lineages. This balance enabled broader genome recovery, including low-abundance populations, and yielded more taxonomically specific results. Despite the slight reduction of the N50 compared to self-coverage binning, this new hybrid binning strategy ensured better grouping of contigs within the same phylum and improved separation between divergent taxa. SR coverage resolution improvement was observed primarily through an increased number of unique and non-redundant bins, which mainly resulted from the addition of new reconstructed bins and the improvement of some bins that were previously retrieved with LR coverage.

In some cases, the number of high-quality bins increased by salvaging contigs and including them in existing bins. SR coverage-based bins involved 113,232 contigs, whereas HiFi-LR coverage allowed the recruitment of only 83,327. Clustering capacity improvement resulted from enhanced coverage depth signal with SR.

A detailed analysis of mid- and high-quality bins further illustrates the impact of SR coverage. For example, with HiFi-LR self-coverage, bin *metabat2*.*163* included 11 contigs, and exhibited a completeness of 93.16% and contamination of 2.98%. When SR coverage was used, 10 of its contigs were regrouped into bin *metabat2*.*1044*, with nearly identical metrics (completeness: 93.02%, contamination: 2.70%). Notably, one contig (ctg3929932_19x_l) was excluded from the SR-coverage bin. This contig had a significantly lower coverage (17X in HiFi-LR, 62X in SR) compared to the mean of the SR bin (92X, standard deviation: 12), suggesting it may have been a coverage outlier. Such redistribution of contigs was likely the cause of the reduced contamination associated with SR-coverage binning (fig. 2B,C). In contrast, the HiFi-LR bin had a broader coverage dispersion (mean: 27X, standard deviation: 10.1), likely causing erroneous contig inclusion. This example highlights how SR coverage refines clustering by improving depth resolution and eliminating poorly supported contigs. The overall taxonomic assignment remained consistent between both bins, assigned to the same lineage within the Burkholderiales order (genus JAABQT01), with nearly identical ANI and tree placement results. Thus, a hybrid binning strategy with SR coverage enhances bin completeness and specificity, stabilising bin composition by filtering out ambiguous contigs, which leads to more taxonomically accurate genome reconstructions.

### Hybrid binning captures a broader soil microbiota diversity

To assign taxonomy to shotgun metagenomic data, we applied two complementary strategies: MAG-based classification using GTDB phylogenomic approach, as described in the previous sections of the manuscript and *in-silico* metabarcoding. *In-silico* metabarcoding refers to identifying and assigning taxonomy to 16S rRNA gene fragments directly extracted from metagenomic reads or contigs, using the SILVA database as a reference[52]. While both methods aim to characterise microbial diversity, MAG-based classification provides more accurate resolution at the genome level, whereas *in-silico* metabarcoding enables direct comparison with taxa obtained by mapping reads (HiFi-LR and SR) on the 16S rRNA reference database. This latter step enables us to evaluate the consistency between taxonomic profiles inferred from metagenomic sequencing and those obtained through standard amplicon-based approaches, which serve as a widely accepted reference in microbial ecology. Additionally, metabarcoding profiles were used as an external benchmark to assess how well the different binning strategies captured the estimated biodiversity in the assemblies.

#### Long-read metagenomics achieves the highest taxonomic overlap with metabarcoding and captures broader microbial diversity

While HiFi-LR sequencing improves assembly contiguity and completeness, SR sequencing provides more informative coverage signals, which are critical for accurate binning. When studying soil samples, a key challenge is to determine how effectively sequencing and binning strategies capture the microbial diversity present in the environment.

To assess this, we compared *in-silico* metabarcoding obtained from metagenomic data with real metabarcoding data obtained from the same soil sample. The metabarcoding approach, targeting the V3-V4 region of the 16S rRNA gene, served here as a proxy for estimation of the microbial diversity in the absence of ground truth and despite biases associated to such data [56]. Taxonomic comparisons were performed based on matches of 16S sequences between metabarcoding data and contigs or reads from the different metagenomic datasets (SR, HiFi-LR, and their corresponding contigs). Given the limited reliability of abundance estimation using metagenomics, these analyses focused solely on the presence/absence of taxa (Fig. 4A).

**Figure 4:**
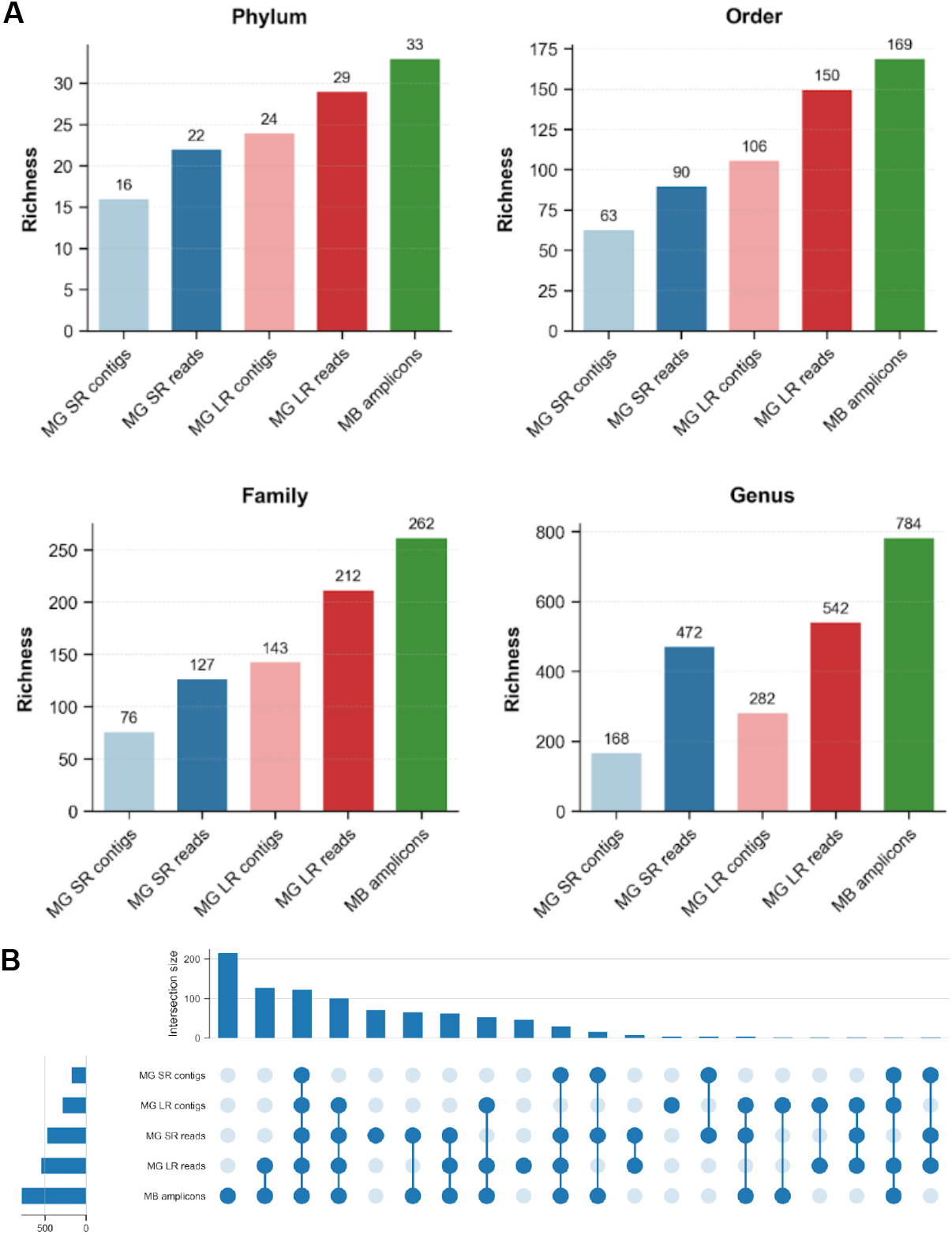
**A)** Comparison of taxonomic richness across sequencing strategies and taxonomic ranks. Bar plots show the number of distinct taxa, richness, identified at four taxonomic levels (Phylum, Order, Family and Genus) using five sequencing approaches: MG SR assembly, MG unassembled SRs, MG HiFi-LR assembly, MG unassembled HiFi-LRs, and MB amplicons, where MG refers to data produced by shotgun metagenomic sequencing and MB: refers to metabarcoding data of 16S V3-V4 region. See also Supp. Table 1. **B)** Taxonomic overlap at the genus level across five datasets: MG SR assembly, MG unassembled SRs, MG HiFi-LR assembly, MG unassembled HiFi-LRs, and MB amplicons, where MG refers to data produced by whole genome metagenomic sequencing and MB refers to metabarcoding data of the 16S V3-V4 region. Horizontal bars on the left of the upset plot represent the total number of genera detected by each method. Vertical bars above indicate the size of each intersection set, corresponding to specific combinations of techniques shown by filled circles below. Overall, the plot illustrates both shared and method-specific taxonomic recovery at the genus level (N = 914). Taxonomic reference confirmed the annotation accuracy improvement through SRcoverage-informed binning of HiFi-LR contigs.

Across all methods, both bacterial and archaeal lineages were consistently identified. However, significant differences emerged when comparing richness at various taxonomic ranks: (i) SR-only metagenomics showed the lowest diversity across most taxonomic levels, except for species (Fig. 4A), where an unusually large number of taxa (7,897 vs. 3,593 in metabarcoding) were detected. This inflation is likely due to low specificity of SR mapping to 16S libraries due to their short length, resulting in overestimation. (ii) HiFi-LR data consistently showed the highest richness among metagenomic datasets and captured taxonomic profiles closer to metabarcoding. At the phylum level, HiFi-LR assemblies and raw reads identified 24 to 29 phyla compared to 33 identified in metabarcoding. (iii) At higher resolution (e.g. genus level), long-read assemblies recovered ∼33% of genera detected by metabarcoding. In contrast, raw HiFi-LR recovered nearly two-thirds of the genera, confirming that the reads forming contigs represent a subset of the total diversity (Table 2, Fig. 4A).

To better visualise taxonomic overlaps, an Upset plot at the genus level (Fig. 4B) illustrates shared and unique genera across the five datasets (SRs, SR assembly, HiFi-LRs, HiFi-LR assembly, and metabarcoding). Metabarcoding recovered the highest number of genera (784), followed by the raw LR dataset. However, only a small subset of genera was shared across all approaches, reflecting detection thresholds and method-specific biases due to the use of databases (i.e., GTDB/SILVA) constructed with different lineage structures. Notably, over 200 genera were uniquely detected by metabarcoding, indicating that a fraction of the microbial community, likely corresponding to rare taxa, remains elusive to metagenomics, even with HiFi-LRs and high depth SRs.

The taxonomic profile provided by metabarcoding sequencing was used as a proxy to assess the taxonomic reliability of reconstructed genomes. We compared GTDB annotations of MAGs with 16S-based annotations of their contigs using the SILVA database (Supp. Fig. 8). Our objective was to assess whether HiFi-LR-based and SR-based coverage at the binning step resulted in differences in MAG quality. Across all taxonomic ranks, annotations from MAG generated using SR coverage were consistently more concordant with SILVA annotations than those from HiFi-LR coverage (Supp. Fig. 8). At the domain level, annotations were consistently assessed across all methods. As the level of resolution increased (from phylum to genera), discrepancies became more frequent, due in part to differences in database content and lineage structure. Despite this, SR-informed binning always yielded a higher number of MAGs with consistent taxonomic placement. Out of 313 bins, 200 MAGs could be compared (due to the absence of the 16S rRNA gene in others), confirming that SR coverage significantly improves both detection and phylogenetic resolution despite persisting discordance with metabarcoding profiles at low taxonomic ranks.

#### HiFi-LR metagenomic microbial detection correlates strongly with taxon abundance

To assess the relationship between microbial abundance and capture efficiency in shotgun metagenomics, we used metabarcoding relative abundance information to compare genera detected and undetected in the HiFi-LR assembly. Figure 5A illustrates the distributions of relative abundances. Genera recovered via metagenomics exhibit significantly higher abundance in the metabarcoding dataset (Wilcoxon test, p = 5.5 × 10^−80^), suggesting a strong positive correlation between microbial abundance and recovery likelihood. Consequently, and as expected metagenomic capture and reconstruction are biased towards dominant taxa. Consistently, the comparison of community profiles across amplicons, bins, and MAGs shows that while dominant phyla such as Actinomycetota, Gemmatimonadota, and Planctomycetota are robustly recovered across approaches, low-abundance phyla (e.g., Chloroflexota, Bacillota) are well represented in amplicon data but tend to be underrepresented in MAGs (Fig. 5B). Importantly, SR coverage information during HiFi-LR binning quantitatively improved the recovery of low-abundance taxa. Specifically, the SR-coverage strategy enabled the reconstruction of 48 MAGs that were not recovered using LR-coverage alone. Taxonomic annotation of these SR-specific MAGs identified 17 phyla, 23 classes, 32 orders, 35 families, and 29 genera, demonstrating that SR-guided binning substantially expands the phylogenetic breadth of reconstructed microbial diversity. This approach effectively reduces the detection gap between metagenomic MAG-identified taxa and metabarcoding profiles.

**Figure 5:**
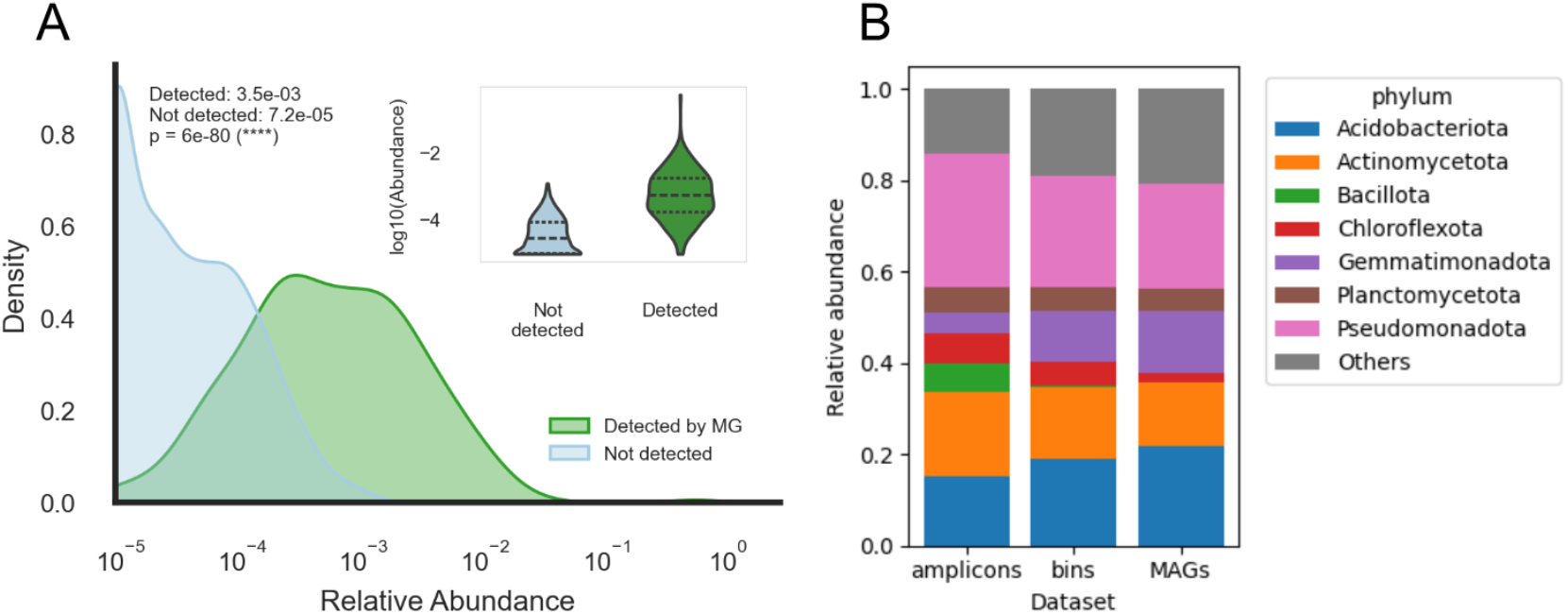
**A)** Distribution of relative abundances for genera assessed by metabarcoding analysis that were either detected or not detected by PacBio HiFi combined metagenomic assembly (Sequel + Revio). Density plots show the relative abundance of genera, classified as either detected (green) or not detected (blue), on a log10 scale using HiFi-LRs. Genera detected by metagenomics (MG) exhibit significantly higher abundances compared to those not detected, with a highly significant difference (Wilcoxon test, p = 6 × 10^−80^). **B)** Relative abundance of phyla among reconstructed bins and MAGs and in the 16S rRNA gene amplicon sequencing dataset (“amplicons”). Phyla accounting for less than 5% of relative abundance were merged into “Others”.

## Discussion

### HiFi-LR sequencing advances genome-resolved soil metagenomics

Soil is among the most microbially diverse environments, with profound implications for global biogeochemical cycles and yet still under-characterised at the genomic level [1, 2]. Although amplicon sequencing and SR metagenomics have provided insight into microbial composition and function, genome-resolved understanding has lagged due to fragmentation, poor resolution of closely related strains, and challenges in assembling repetitive regions [11, 15]. Recent advances in long-read technologies—particularly PacBio HiFi sequencing— have demonstrated their potential in the gut microbiome, enabling highly contiguous assemblies and more comprehensive recovery of the gene sets [18, 44, 57]. Here, we explore their benefits in a soil microbiome sample, directly comparing the added value of several sequencing and binning strategies. We show that HiFi-LRs substantially improve assembly quality as well as MAG completeness and contiguity compared to Illumina SRs alone, even at moderate sequencing depth. This improvement is expected to support more precise taxonomic and functional annotation in future studies, providing a robust foundation for investigating microbe–microbe and microbe–environment interactions in soil.

### Hybrid binning of HiFi-LR contigs with SR data enhances recovery and resolution of MAGs

Despite the benefits of long reads, MAG recovery from long-read assemblies can be constrained by low coverage and weak abundance signals, particularly for rare taxa. In our work, coverage estimates from HiFi-LR data alone were poorly effective for abundance-based binning, even though an entire Sequel II and entire Revio SMRT Cell were used for sequencing of a unique sample. This low coverage signal resulted in contamination and collapsed bins. To address these challenges, we used the high-coverage Illumina SR data to generate coverage profiles for HiFi-assembled contigs. This hybrid binning strategy significantly improved bin quality—reducing contamination, increasing dereplication, and resulting in a 24% increase in MAG yield. Notably, this approach enabled the recovery of additional MAGs from low-abundance and taxonomically underrepresented lineages, including 34 genera not detected with other strategies and several lacking close matches in reference databases. We also observed improved resolution in phyla such as Acidobacteriota and Planctomycetota, consistent with previous successes of hybrid methods in less complex microbiomes [16]. We note, however, that MAG counts alone are not a comprehensive quality measure and that bin segmentation, strain heterogeneity and contig taxonomic discordance within bins are additional valuable metrics to consider.

By testing the impact of subsampling the SR dataset used for coverage estimation during binning, we showed that SR sequencing depth is a critical factor for improving MAG recovery. This indicates that binning may remain a major bottleneck in LR-only strategies, particularly as long-read throughput is still limited compared to SR. We also explored alternative hybrid strategies, such as combining SR and LR data at the assembly step [16, 22, 23]. However, the size and complexity of our soil dataset posed significant challenges for current hybrid assemblers, often requiring extensive manual intervention or failing to complete successfully despite substantial computational resources. Although partial hybrid assemblies improved read recruitment, these gains did not translate into improved MAG quality in our dataset. This suggests that, for highly complex soil microbiomes, using SR data for coverage-guided binning of LR assemblies is presently a more scalable and robust approach than full hybrid metagenomic assembly, while still capturing most of the complementary information provided by short reads.

Datasets integrating multiple sequencing platforms, and in particular HiFi long reads, remain scarce (49–51). Our dataset—including 81 Gbp of PacBio HiFi reads and 147 Gbp of Illumina ultra deep sequencing short reads from the same soil sample, as well as 16S rRNA gene amplicon data—provides a valuable benchmark for soil metagenomics.

### Limitations and future perspectives

Despite the improvements offered by HiFi-LR sequencing and hybrid binning, the number of recovered MAGs remains substantially lower than the taxonomic richness estimated by amplicon-based profiling. For complex communities such as soil, the absence of a ground truth makes comprehensive benchmarking challenging. Current standards rely on mock or synthetic communities [58], which do not fully capture environmental complexity. In this study, we use metabarcoding data as a proxy to estimate an upper bound for microbial diversity, while acknowledging the known limitations of this approach, including amplification bias, copy number variation, and taxonomic assignment inaccuracies [56]. The discrepancy between 16S rRNA gene-based estimates and the number of recovered MAGs has been frequently reported [7, 12], and is likely even greater in soil due to its extreme heterogeneity and prevalence of low-abundance or recalcitrant taxa. It is also important to note that our analyses were performed on a single soil sample, limiting the extent to which these findings can be generalized to other soils or environmental contexts.

Another limitation is the cost currently associated with HiFi-LR sequencing. While increasing sequencing depth—potentially with multiple SMRT Cells—could improve recovery of genomes from low-abundance species, this was not evaluated in our study and remains financially prohibitive for many projects. Thus, cost remains a significant factor that may restrict large-scale adoption of this approach. We also note that integrating data from two sequencing platforms (PacBio and Illumina) introduces some additional steps compared to single-platform workflows. However, the bioinformatics tools and approaches employed are state-of-the-art and widely used in metagenomic studies, making the workflow re-usable for most research groups with standard resources and expertise.

A technical constraint in soil metagenomics is the relatively short length of extracted DNA, rarely exceeding 6 kb, which is well below the 30–40 kb typically obtained from gut or aquatic samples [59]. This limits the full potential of HiFi and other long-read technologies. Thus, the development of improved DNA extraction methods that minimize shearing and efficiently remove inhibitors is of importance. Achieving greater fragment lengths will enhance assembly contiguity and enable full-length resolution of repeats, further improving MAG quality and yield.

Looking forward, advances in high-throughput HiFi sequencing, optimised protocols for extracting long DNA fragments from challenging samples, and continued expansion of environmental genome catalogs will be essential. Together, these improvements will enable more comprehensive and functionally resolved characterisation of soil microbial communities, contributing to our understanding of their roles in nutrient cycling, soil health, and ecosystem resilience.

## Conclusion

Collectively, our results demonstrate that integrating PacBio HiFi-LR sequencing with SR coverage for binning markedly enhances the recovery, resolution, and diversity of MAGs from complex soil environments. This hybrid strategy represents a step toward closing the gap between microbial diversity estimated with amplicon-based profiling and that captured by shotgun metagenomic assembly, facilitating access to previously undetected lineages and enabling a more comprehensive taxonomic characterization of soil microbiomes. By leveraging the high contiguity of long reads and the depth provided by short reads, the hybrid approach improves binning accuracy and supports the recovery of a broader range of genomes, including those from low-abundance taxa.

Importantly, while HiFi-LR sequencing alone offers advantages in assembly quality, the addition of short-read data for binning substantially increases the resolution and completeness of recovered MAGs. Our findings underscore the potential of this hybrid method both for enhancing new studies and, whenever samples are still available, for adding value to existing SR datasets. However, the benefits of this approach are contingent on adequate sequencing depth, which remains a limiting factor, particularly for the recovery of rare taxa in highly complex soil microbiomes.

In summary, the integration of HiFi-LR and SR sequencing represents a robust and scalable strategy for genome-resolved exploration of complex microbial ecosystems. As sequencing technologies and library preparation methods continue to advance, and as comprehensive reference catalogues expand, the potential for soil metagenomics to reveal the functional roles and interactions within microbial communities will continue to grow.

## Supporting information

SuppData

## Declarations

### Availability of data and material

All data generated or analysed during this study are publicly available. The datasets and custom scripts have been deposited in the *Recherche Data Gouv* repository under the Soil Metagenome Binning collection [60].

Sequencing and assembly data are available through the European Bioinformatics Institute (EBI) under the following accession number: PRJEB97182.

### Ethics approval and consent to participate

Not applicable.

### Consent for publication

Not applicable.

### Authors’ contributions

C.B., N.M., S.M., A.F., M.B.-B., C.L., J.-M.F., C.F., and E.G.J.D. conceived and designed the study.

C.B., N.M., A.P. developed and implemented the software and methodological approaches.

C.B., N.M., and A.F. carried out the analyses and formal data interpretation.

C.B. curated the data.

C.B., N.M., C.F., and É.G.J.D. prepared the original draft of the manuscript.

C.L., R.V., C.F., and É.G.J.D. supervised the project.

M.B.-B., A.F., P.A., D.J.S., C.F., and É.G.J.D. acquired funding.

All authors reviewed and approved the final manuscript.

## Acknowledgement

We sincerely thank the Sophia Agrobiotech bioinformatics platform for its valuable support and expertise in data processing, as well as for providing optimised computing hardware and software [61]. We also thank Genotoul for providing high computing resources and fast reactivity during access openings [62]. We acknowledge the GenOuest bioinformatics core facility (https://www.genouest.org) for providing the computing infrastructure. Some experiments presented in this paper were also carried out using the PlaFRIM experimental testbed, supported by Inria, CNRS (LABRI and IMB), Université de Bordeaux, Bordeaux INP and Conseil Régional d’Aquitaine (see https://www.plafrim.fr).

Part of the experiments (short read metagenomic sequencing, 16S rRNA gene sequencing) were performed at the PGTB platform [63]. We are grateful to the Genosol, Gentyane, Get-Plage and PGTB platforms for their significant contributions, including soil DNA extraction and high-throughput sequencing, which have been essential for the progress of this work. We are grateful to all collaborators and technical staff who have contributed their expertise and assistance throughout this research. This work benefited from the use of the GenoSol-CRB and GenoSol-LADM services of the GenoSol platform [64], hosted by UMR 1347 Agroecology at the INRAE Bourgogne–Franche-Comté center. GenoSol-CRB is part of BRC4Env [65], while GenoSol-LADM is integrated into the AnaEE-France research infrastructure. This work was performed in collaboration with the GeT core facility, Toulouse, France [66]. We sincerely thank Isabelle Kupin for her assistance in depositing the datasets.

