## Supplementary material for "Accurate MAG reconstruction from complex soil microbiome through combined short- and HiFi long-reads metagenomics": SuppData

### Appendices

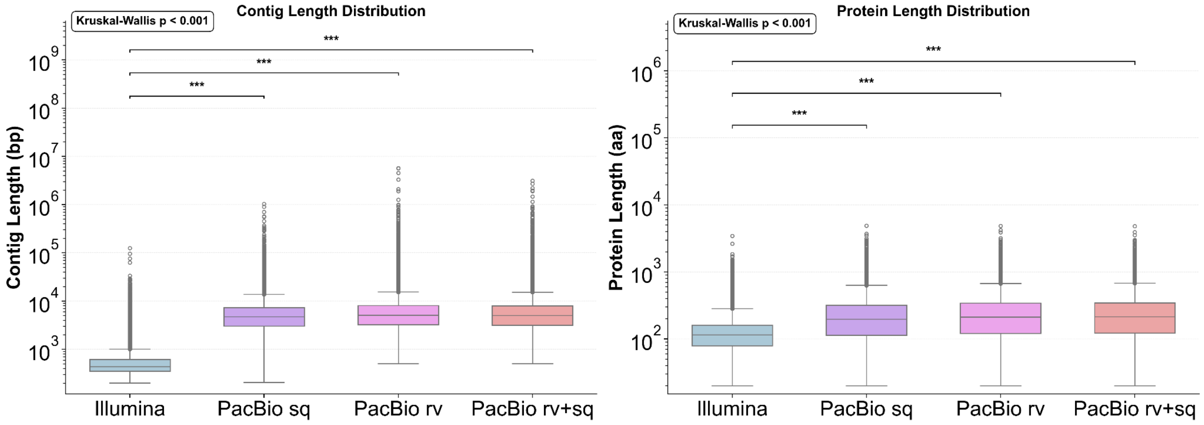


Figure 1. Comparison of contig (bp) and protein length (aa) distributions across sequencing technologies, PacBio Sequel (sq), PacBio Revio (rv), and PacBio combined (rv+sq). Long-read assemblies (PacBio-based) yield significantly longer contigs and protein-coding sequences compared to Illumina assemblies (Kruskal–Wallis test, $p < 0.001$; *** $p < 0.001$).

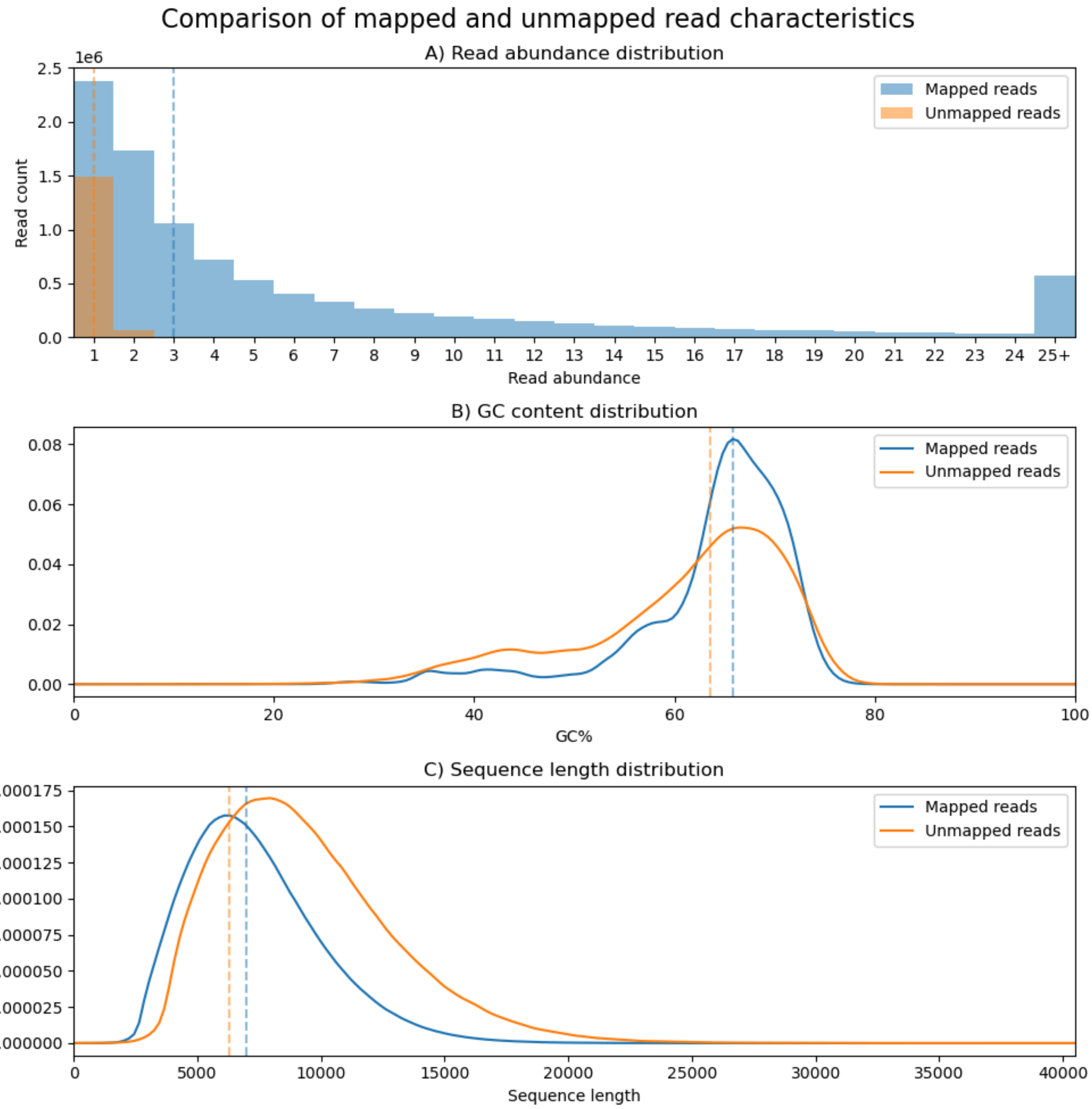


Figure 2. Comparison of mapped and unmapped reads characteristics within the sequel + revio long reads dataset. Shown in dotted lines are the median values. Read abundance distribution (A), is calculated as the median occurrence (within the full reads dataset) of the 27-mers within that read.

Table 2. Comparative analysis of Richness across natural soil environmental samples using Metabarcoding (MB) and Metagenomics (MG) approaches for the V3-V4 region of 16S rRNA markers reported for four foremost taxonomic ranks.

| Taxa rank | MG SR assembly | Mg SR reads | MG LR assembly | MG LR reads | MB V3-V4 |
|---|---|---|---|---|---|
| kingdom | 2 | 2 | 2 | 2 | 2 |
| phylum | 16 | 22 | 24 | 29 | 33 |
| order | 63 | 90 | 106 | 150 | 169 |
| family | 76 | 127 | 143 | 212 | 262 |
| genus | 168 | 472 | 282 | 542 | 784 |
| species | 1791 | 7897 | 352 | 1109 | 3593 |

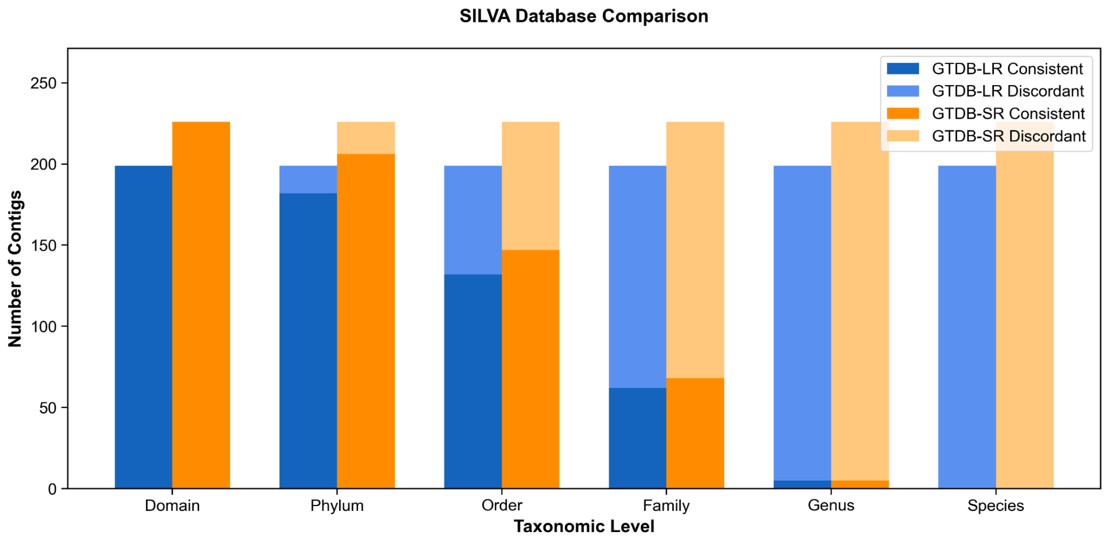


**Figure 3** – Taxonomic annotation of MAGs using GTDB-tk and 16S detection against the SILVA database.

Comparison of GTDB-tk and SILVA-derived annotations across ranks. Stacked bar plots display the number of contigs annotated at each taxonomic level (from domain to species) for LR and SR assemblies. Consistent (dark shades) and discordant (light shades) assignments concerning SILVA are shown for each GTDB-tk assignment, highlighting agreement and divergence across taxonomic assignment approaches.
